# Innocent until proven guilty: Privacy-preserving search over a central CODIS criminal database from the field

**DOI:** 10.1101/2020.06.22.164095

**Authors:** Jacob A. Blindenbach, Karthik A. Jagadeesh, Gill Bejerano, David J. Wu

**Affiliations:** Department of Computer Science, University of Virginia, Charlottesville, VA; Klarman Cell Observatory, Broad Institute of MIT and Harvard, Cambridge, MA; Department of Computer Science, Stanford University, Stanford, CA; Department of Developmental Biology, Stanford University, Stanford, CA; Department of Pediatrics (Medical Genetics), Stanford University, Stanford, CA; Department of Biomedical Data Science, Stanford University, Stanford, CA

## Abstract

The presumption of innocence (i.e., the principle that one is considered innocent until proven guilty) is a cornerstone of the criminal justice system in many countries, including the United States. DNA analysis is an important tool for criminal investigations^1^. In the U.S. alone, it has already aided in over half a million investigations using the Combined DNA Index System (CODIS) and associated DNA databases^2^. CODIS includes DNA profiles of crime scene forensic samples, convicted offenders, missing persons and more. The CODIS framework is currently used by over 50 other countries^3^ including much of Europe, Canada, China and more. During investigations, DNA samples can be collected from multiple individuals who may have had access to, or were found near a crime scene, in the hope of finding a single criminal match^4^. Controversially, CODIS samples are sometimes retained from adults and juveniles despite *not* yielding any database match^4–6^. Here we introduce a cryptographic algorithm that finds any and all matches of a person’s DNA profile against a CODIS database *without* revealing anything about the person’s profile to the database provider. With our protocol, matches are immediately identified as before; however, individuals who do *not* match anything in the database retain their full privacy. Our novel algorithm runs in 40 seconds on a CODIS database of 1,000,000 entries, enabling its use to privately screen potentially-innocent suspects even in the field.

## Introduction

DNA-based forensic analysis is a powerful tool used by law enforcement agencies around the world for solving crimes^1–3^. With today’s technology, local police stations^7^ and even agents in the field^8^ can generate a suspect’s DNA profile to search against central criminal DNA databases in an impressive 90 minutes^9,10^. At the same time, the increased prominence of DNA-based forensics opens up new avenues for misuse including social control and racial profiling^11,12^.

In particular, the storage of DNA samples from potentially innocent individuals *permanently* links their genetic identities to criminal databases (without due process). Within the United States, these controversial practices have included collecting DNA from individuals who have been arrested but not convicted or even charged with a crime^4^, people who are not even arrested (so-called “stop-and-spit” and “swab-and-go” practices^13^), detained immigrants and asylum seekers^6^, and even children brought in for questioning^5^.

Here we develop a novel solution (Figure 1) whereby an agent in the field, using modest computational resources, could privately query a suspect’s DNA profile against a large central database of DNA profiles. In less than 40 seconds, the agent learns whether the suspect’s profile matches, based on CODIS rules, against any profile in the central database of 1,000,000 profiles, while learning nothing else about the contents of the central database. More importantly, the central database itself learns absolutely *nothing* about the DNA profile being searched. Any match discovered can be investigated further as before. However, should no match be made, the suspect’s DNA profile, now exonerated, can be disposed of on the spot, with zero risk that the central database provider chose to retain it.

**Figure 1.**
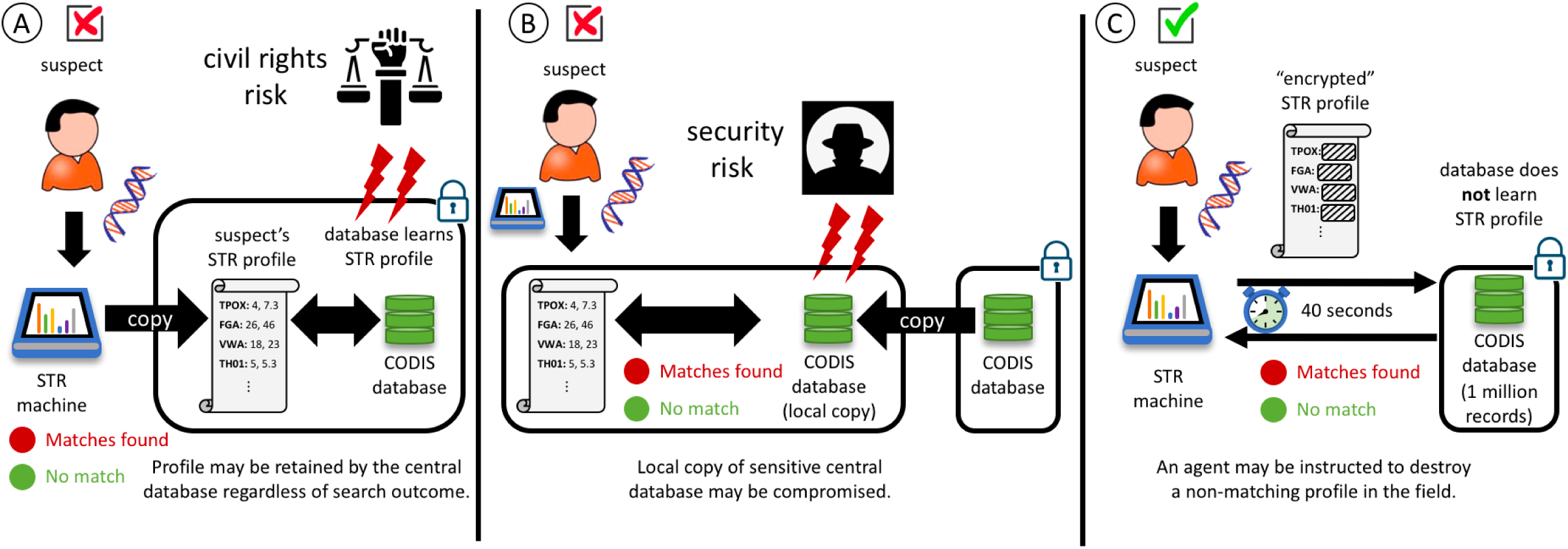
A novel privacy-preserving CODIS DNA profile matching protocol. Rapid STR profiling technologies enables genetic testing and matching from the field. They provide a valuable tool for crime solving but raise significant civil rights concerns regarding data retention and racial profiling. (A) Today, STR profiles collected from potentially innocent individuals are sent to a central CODIS database to check for matches. Here, the central database learns the full query profile and has the option to retain it, irrespective of whether the search yields any match or not. (B) To try and provide anonymity to exonerated (unmatched) profiles, one may load a private copy of the central database onto every field device. This way, an unmatched suspect profile may be destroyed in the field to retain suspect privacy. However, this approach risks exposure of all or parts of the sensitive central database to malicious parties who get ahold of a field device. (C) Our privacy-preserving search protocol enables a new approach where agents can still query a central CODIS database as in (A), but in a way that completely hides the query from the central database. The agent still learns the outcome of the query as before. However, an innocent profile, not matching anything in the database may safely be destroyed in the field. The central database can no longer store it, as it has learned nothing about it through the query.

## Results

### CODIS system of DNA profiles and profile matches

The United States CODIS system originally established a set of 13 loci across the genome coinciding with short tandem repeats (STRs) as a method for comparing genetic data for identification purposes^3^. These were expanded to a set of 20 loci in 2017. Many countries have adopted a similar system with a combination of existing core STR loci and region-specific STR loci. For example, a European Union (EU) system of 16 loci, a UK system of 11 loci and a Chinese system of 20 loci have all been described^14,15^ (see Online Methods and Supplementary Table 1). At each locus, individuals have 2 alleles, one inherited (with possible personal modification) from each parent. The allele at each locus is represented by a varying range and represents the number of repeats of a 2 to 6-character (base pair) generic sequence observed in the individual’s genome. Based on frequency statistics collected by the FBI, the probability that two unrelated individuals share the same STR profile across all 13 core loci positions of the older US system is 1 in 575 trillion^16^. This probability further decreases with the expanded set of 20 loci introduced in 2017.

The CODIS system describes several ways to query a database of STR profiles. The standard and default method is a “high-stringency search with one mismatch,” which requires that both of the alleles appearing in at least 19 out of the 20 STR loci between the query profile and the database profile match exactly^17,18^. Central CODIS databases holding thousands to millions of DNA profiles that can be used for such queries are maintained in the US at the national, state, and sometimes even municipal level^2^ (as well as very similar databases and matching rules in dozens of other countries^3^).

### Encoding an STR profile for secure computation

We encode an STR profile as a binary string. For each STR locus, we propose a public dictionary that maps every pair of STR alleles to a unique binary string. The number of bits used to encode each allele is determined based on the number of unique alleles expected at the locus^14^. We assume that all entries in the central database (unknown to the agent making the query) and the suspect’s STR profile held by the agent (unknown to the central database) are encoded in this commonly-agreed upon manner.

### Secure protocol overview

It suffices to construct a secure protocol for comparing a single entry in the central database with the suspect’s profile. By running this protocol over and over against all entries in the central database, the agent will learn the indices of the complete set of matching records, and nothing else, while the central database will learn nothing (Figure 1C).

We start by fitting a computational model to the task, before later securing it: given two binary strings encoding two STR profiles as above, decide whether they correspond to a match according to the CODIS specification or not. Our key observation is that one can efficiently compute this using a compact deterministic finite automaton^19^ (DFA). In a DFA, the computation begins at an initial state, and at each step of the computation, the DFA reads a bit of the input and advances the state. After reading all of the input bits, the DFA ends in either an “accepting” state or a “rejecting” state. For example, a DFA can be constructed to test for equality between two equal length bit-strings by defining two sets of states: a set of “matching” states and another for “mismatching” states. The computation begins in the “matching” set, and as each bit of the input is read, it is compared against the target bit. As long as the current state is in the matching set, if the two bits match, the program transitions to the next state in the matching set, and otherwise, it transitions to a state in the mismatching set. Once a single bit mismatches, one enters the mismatching set, from which all inputs lead only to the next state in the mismatching set. The input bit string is equal to the target string if the computation concludes in a state in the matching set, and is otherwise unequal (see Supplementary Figure 1). We use this DFA to check for matches at a single STR locus. A similar DFA can be used to test for equality with up to one mismatch (see Online Methods and Supplementary Figure 2). We use this one to compute a CODIS match of at least 19 of 20 STR loci.

In our setting, the central database owner constructs a DFA for each profile in its database. The input to the DFA is the agent-held suspect’s STR profile. The DFA computation ends in an accepting state if the suspect’s profile is a CODIS match to the central database profile; otherwise, the DFA computation ends in a rejecting state.

Our protocol now proceeds as follows. At the beginning, the agent knows the initial state of the DFA as well as the suspect’s STR profile (hidden from the central database). The central database holds the DFA corresponding to a database entry (hidden from the agent). Using a cryptographic protocol called “oblivious transfer^20,21^,” the agent and the central database now jointly perform the evaluation of the DFA on the agent’s input STR profile.

Specifically, at each step of the DFA evaluation, the central database enumerates *all* possible states of the DFA that the agent might be in and all possible states the agent will end up in based on the next bit of the agent’s input. These statements are of the form “if you are in state *X* and the next bit of your input is 0, then you will proceed to state *Y*.” Using the oblivious transfer protocol, the agent can secretly choose to learn exactly one of these statements *without* revealing which one she chose to the database server. In this case, the agent chooses the statement corresponding to her current state and the next bit of her input; this in turns reveals to the agent her next state in the DFA evaluation. The oblivious transfer protocol hides all of the other statements from the agent, so the agent cannot learn what would have happened had she been in a different state or had a different input.

At the very end of the protocol, the agent arrives at either an accepting state or a rejecting state, which indicates whether the STR profile she provided matched against the central database’s profile or not. As described so far, the agent learns the full execution path in the DFA as well as whether her input profile matches the database profile or not. To ensure that the execution path does not leak additional information about the profiles on the database server, the central database additionally encrypts all of the intermediate steps of the computation (in a way that still enables the above evaluation procedure). Then, at each of the intermediate steps, the agent no longer knows where she is in the actual DFA execution. The central database only provides a single intelligible state: the very last state which reveals whether the two profiles match or not match.

By repeating the above protocol with each profile in the central database, while changing the encryption of intermediate states every run, the agent has learned just one thing: the database indices of any and all profile matches. She learns nothing else about any of the centrally-stored profiles. On the flip side, the central database learns nothing at all about the suspect’s STR profile; at every step, they only provide an exhaustive list of “if you are here, and have this bit next, then go there.” We have thus achieved the desired goal of Figure 1C. We encourage our readers to refer to the Online Methods for the full technical details and security analysis.

### Performance measurements

With our protocol implementation, an agent in the field in Northern California can privately query a CODIS database containing 1 million STR records located 3,000 miles away in Northern Virginia in just 38 seconds, using 180 MB of online communication. In our experiments, we represent each STR profile as a vector of 20 biallelic components (212 bits in total) based on the current US CODIS specification^2^. Our protocol additionally requires preparing 116 million oblivious transfer correlations^20,21^ and 122 MB of client-side storage (see Online Methods and Supplementary Table 2). These can be generated in a separate preprocessing phase on commodity hardware in about a minute using existing state-of-the-art oblivious transfer extension protocols^22^. Since these correlations are independent of both the query and the database contents, they can be prepared concurrently with the 90 minutes needed for STR profile derivation^9,10^, and thus, contributes no extra latency.

We also measured the performance of our protocol on CODIS specifications reported for the UK^14^, the EU^14,18^, and China^15^ (see Online Methods for specifications details). In all cases, the cost of the protocol is smaller or comparable to that of the US CODIS system (up to 40 seconds and under 200 MB of communication; see Table 1). To illustrate the scalability of our approach, we also measured the performance for a CODIS system with 40 loci and an encoding length of 492 bits, a setting based on a system previously tested by NIST^23^. Performing a CODIS search in this setting over a database of a million records completes in just 72 seconds and requires only 340 MB of communication. Thus, our protocol also scales favorably to future scenarios with an expanded set of STR loci.

We also tested the performance of our protocol on central databases of different sizes (Figure 2). Both the online bandwidth and execution time of our protocol scale linearly with the size of the STR profile (a function of both the number of loci and the number of bits needed to represent the allele values at each locus) and with the size of the central database. For example, our protocol can search over a 20-STR U.S. database with 10 million records in just under 6 minutes.

**Figure 2.**
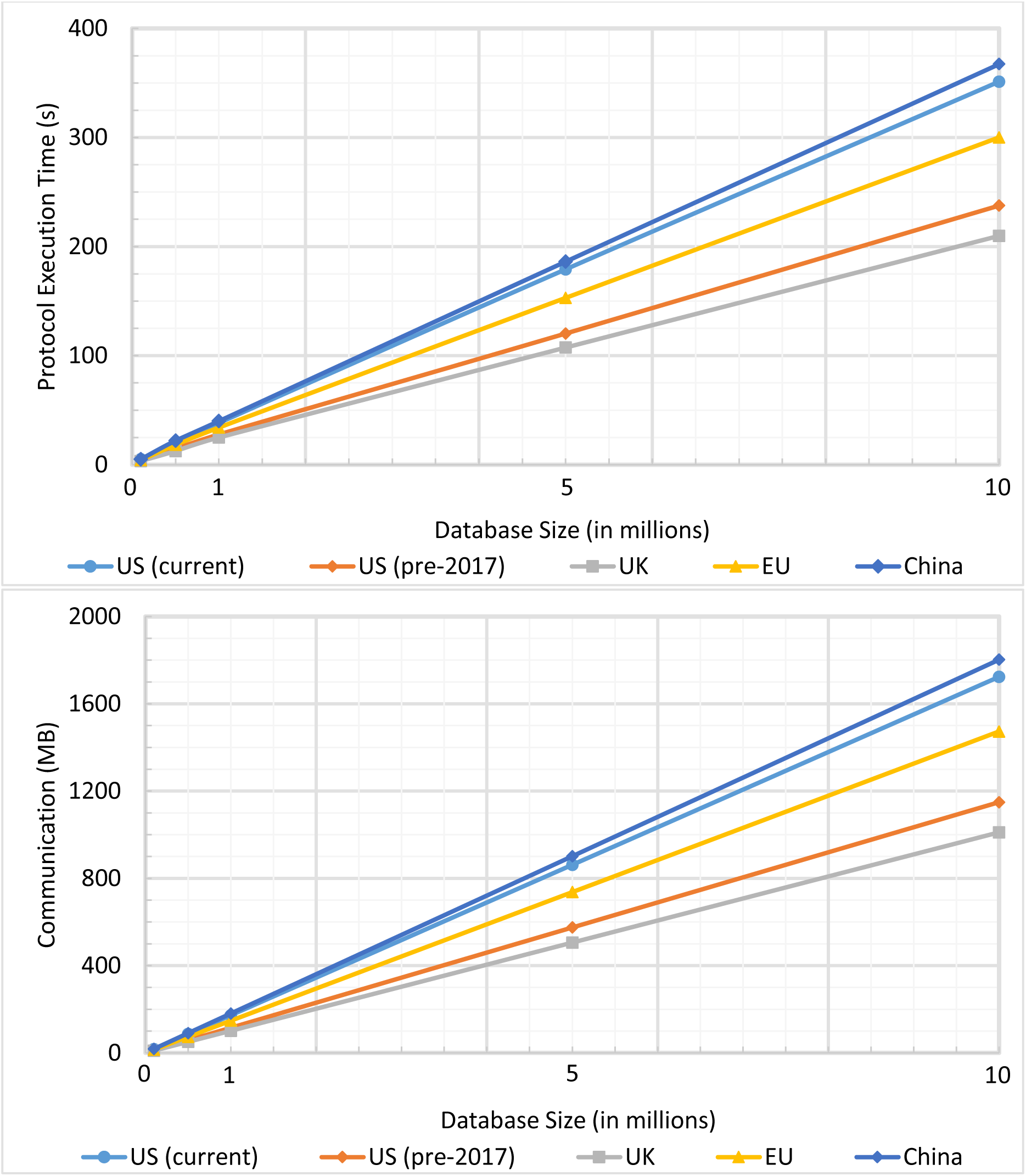
Single query search performance against an entire CODIS database as a function of database size. The number of STR loci and number of precision bits assumed for each system are described in Supplementary Table 1.

## Discussion

DNA profiles are an extremely powerful tool in forensics and crime solving^24^. Our law enforcement agencies have a duty to both serve and protect their communities. How can they use DNA databases to find criminals while simultaneously protecting the privacy of innocent individuals who make up the majority of each society^12^? It is natural to think that searching a suspect’s DNA profile against a master database either requires the database to see the suspect’s profile, or for the agent holding the suspect profile to have a local copy of the master database. Both scenarios compromise privacy (Figure 1). Here we show a third way, whereby a field agent searches the remote master database without learning anything about it except any possible match. Moreover, the central database aids the search while learning *nothing* about the suspect’s profile. Should the search come up empty, if the field agent disposes of the sample, they have both served their community and protected its privacy.

DNA profiling machines can now routinely produce a searchable profile in a matter of 90 minutes, and can even be carried to the field^8^. Integrating our privacy-preserving protocol with such a system adds minimal overhead and can easily fit into existing workflows. In fact, the computational requirements to run the protocol are so modest, it is likely they can be performed on a modern smartphone. The performance of our protocol also compares favorably against generic approaches for privacy-preserving computation. For instance, a direct implementation of a query protocol using Yao’s garbled circuits^25^ would require communicating, storing, and evaluating a circuit that is around 8 GB (*≈* 260 million AND gates) in size for a database with 1 million profiles (and increase to 80 GB of communication and storage for a 10 million entry database). In comparison, our protocol requires less than 125 MB of offline storage for the OT correlations (which can be generated in about a minute^22^) for a database of 1 million profiles.

Our implementation follows the CODIS guidelines for high-stringency matching (the default mode of searching)^17,18^.The deterministic finite automata at the root of our approach can be easily extended to also support moderate and low-stringency matches as well as partial match queries, with modest increases in computation time and communication. In fact, similar operations like paternity testing and ethnicity identification can also be formulated as a similar string-matching problem and implemented using a similar approach. In all cases, the correct answer is obtained while the input DNA profiles to the computation remain private. Our protocol is relevant not only in the US, but also in any of dozens of countries that use a CODIS-like system^3^. It scales well with the size of the central database (Figure 2), on current hardware that will only get faster, and most importantly, it gives the agent in the field or local office, the ability to destroy an exonerated profile that has yielded no incriminating match. Whether they are instructed to do so or not is a civil rights matter that each country must resolve for itself^11,12,26,27^. The importance of our work is in showing that accurate practical implementations to enable these fundamental rights are already doable.

## Online Methods

### STR profiles and representation

The US National Institute of Standards and Technology (NIST) has enumerated all possible allele values for each STR core locus in the current US system^14^. While over 50 other countries reportedly use a local variant of the same CODIS system^3^, details of each country’s system, or even a list of these 50+ countries are not easily found. Possibly current versions of the EU and UK systems are described by NIST^14^, while a partial description of the Chinese system is provided in a paper co-authored by officials from the Chinese Ministry of Justice^15^. Similarly, the high-stringency matching rule with at most one mismatching locus is published for the US^17^ and EU^18^. As the exact CODIS system parameters are immaterial for the essence of our proposed solution, when necessary, we infer the number of possible values at a locus based on the existing NIST standards, and also assume a default search configuration of high-stringency matching of all but one locus for all systems.

In 2017, the US 13 loci system was replaced by the 20 loci system^3^. It is unclear if or when this set will be expanded on. For purposes of illustrating the scalability of our solution we use a 40 loci system described in a NIST paper partly funded by the FBI^23^.

### Security model

We work in a two-party setting where the server holds a profile ***v*** = (*v*_1_, *…, v*_*n*_) and the client holds a query vector ***w*** = (*w*_1_, *…, w*_*n*_), where each of the components *v*_*i*_, *w*_*i*_ ∈ {0,1}^*𝓁*^ can be represented by *𝓁*-bit strings. We define the “threshold matching” function TM_*k*_(***v, w***) to be the following Boolean-valued function:

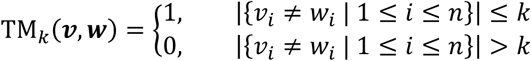

In words, TM_*k*_(***v, w***) outputs 1 if the vectors ***v*** and ***w*** disagree on at most *k* components. In this work, we design a secure protocol such that at the end of the protocol, the client learns the value of TM_*k*_(***v, w***) and nothing more about the server’s profile ***v***, while the server learns nothing about the client’s query ***w***. Importantly, the client only learns whether the number of differing components between ***v*** and ***w*** is greater than *k* or not, but nothing about the exact number of differing components or the indices of the differing components.

We can naturally extend this notion to the setting where the server holds a database with many vectors {***v***_1_, *…*, ***v***_*t*_} and the client’s goal is to learn all of the indices *i* ∈ {1, *…, t*} where TM_*k*_(***v***_*i*_, ***w***) = 1, but nothing more about the vectors ***v***_1_, *…*, ***v***_*t*_. Since a protocol for private evaluation of TM_*k*_(***v, w***) for a single entry ***v*** suffices for searching over a database of values (by repeating the protocol for each database entry ***v***_1_, *…*, ***v***_*t*_), we focus on the single-instance setting for the remainder of this section.

Throughout this work, we assume that the client and the server are semi-honest or “honest-but-curious;” namely, both the client and the server will follow the protocol as described, but may try to infer additional information about the other party’s private inputs based on the messages they receive in the protocol. We note that our protocol actually ensures privacy of the client’s query even against a *malicious* database server that may arbitrarily deviate from the protocol execution, provided that the underlying “oblivious transfer” protocol we use (see below) is secure against a malicious sender.

### Oblivious transfer and the OT-hybrid model

An oblivious transfer (OT) protocol^20,21,28^ is a two-party protocol between a sender and a receiver. In a 1-out-of-*k* OT protocol on *t*-bit messages, the sender holds *k* messages *m*_1_, *…, m*_*k*_ ∈ {0,1}^*t*^ while the receiver holds an index *i* ∈ {1, *…, k*}. At the end of the protocol, the receiver learns the *i*^th^ message *m*_*i*_ and nothing else about any of the remaining messages, while the sender learns nothing. To facilitate the analysis of our protocol, we will work in the “OT-hybrid” model where we assume that the parties have access to a *trusted party* that implements the above 1-out-of-*k* OT functionality^20^. We can then replace the trusted party with a cryptographic implementation of a 1-out-of-*k* OT protocol^28^. If our protocol provides semi-honest security in the OT-hybrid model, and we instantiate the OT protocol with a cryptographic protocol that is secure against semi-honest adversaries, then the overall protocol is also secure against semi-honest adversaries (*without* relying on any trusted party)^29^.

### Oblivious transfer correlations

We can significantly reduce the *online* cost of oblivious transfer by first precomputing *input-independent* “oblivious transfer correlations” in an offline (or preprocessing phase). Because the correlations are input-independent, they can be precomputed without knowledge of the client’s query or the server’s database. These OT correlations can be generated efficiently using a technique called OT extension in a separate input-independent preprocessing step^30^ (this can even be done with low communication using a recent approach called silent OT extension^31^). Alternatively, they can be generated ahead of time by a trusted dealer or a secure hardware platform (observe that in both of these settings, the party generating the correlations does *not* need to know anything about the query or the database entries).

Very briefly, a 1-out-of-*k* OT correlation for *t*-bit messages consists of the following: (1) a tuple of *k* random values *r*_1_, *…, r*_*k*_ ∈ {0,1}^*t*^ for the server; and (2) a random index *β* ∈ {1, *…, k*} together with the value *r*_*β*_ for the receiver. Once the client and server have this OT correlation, it is straightforward to implement a 2-message 1-out-of-*k* OT (on an arbitrary collection of sender messages *m*_1_, *…, m*_*k*_ ∈ {0,1}^*t*^ and receiver index *i* ∈ {1, *…, k*}). We recall the construction below:

- **Receiver message:** The receiver sends the index *j* = *i* + *β* (mod *k*) to the sender. Observe that since *β* is uniformly random over {1, *…, k*} and unknown to the sender, this message perfectly hides the receiver’s index *i*. For notational convenience, we will consider the output of arithmetic modulo *k* to be a value between 1 and *k* (as opposed to 0 and *k* − 1).
- **Sender response:** On input an index *j* ∈ {1, *…, k*} from the receiver, the sender computes the blinded message *c*_*i*_ ← *m*_*i*_ ⊕ *r*_*j*−*i* (mod *k*)_, where ⊕ denotes the bitwise exclusive-or operator (i.e., bitwise xor). The sender sends the blinded messages *c*_1_, *…, c*_*k*_ to the receiver. Since *r*_1_, *…, r*_*k*_ are uniform over {0,1}^*t*^ and the receiver knows exactly one of these values (i.e., *r*_*β*_), *k* − 1 out of the *k* values are perfectly hidden from the receiver.
- **Receiver reconstruction:** The receiver computes its message as *c*_*i*_ ⊕ *r*_*β*_.

Correctness of the protocol follows from the following simple relation:

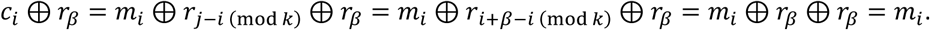

To summarize, a 1-out-of-*k* OT correlation on *t*-bit values yields a 1-out-of-*k* OT on *t*-bit messages in two rounds of interaction where the total communication is ⌈log *k*⌉ + *kt* bits. The only necessary computation is integer arithmetic and bitwise operations, so this is a very lightweight protocol.

### Representing threshold matching as a deterministic finite automaton

Given two vectors ***v*** = (*v*_1_, *…, v*_*n*_) and ***w*** = (*w*_1_, *…, w*_*n*_) where *v*_*i*_, *w*_*i*_ ∈ {0,1}^*𝓁*^, our objective is to securely compute the threshold matching function TM_*k*_(***v, w***) defined above. In our setting, we assume the database server holds ***v*** while the client holds ***w***. The starting point of our design is to express the computation of TM_*k*_ as a composition of two deterministic finite automatons (DFAs):

- The first DFA checks equality of *𝓁*-bit strings. Namely, for a string *v* ∈ {0,1}^*𝓁*^, the machine computes the function *g*_*v*_(*w*) that outputs 1 if *v* = *w* and 0 otherwise.
- The second DFA computes a threshold function. Namely, for a target sequence of bits *a*_1_, *…, a*_*n*_ ∈ {0,1} and a threshold *k*, the machine computes the function 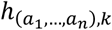 (*b*_1_, *…, b*_*n*_) that outputs 1 if *a*_*i*_ ≠ *b*_*i*_ on at most *k* indices 1 ≤ *i* ≤ *n* and 0 otherwise.

**Table 1.** Runtime and network communication needed to privately compare a suspect CODIS profile in the field to a central office database far away. End-to-end protocol execution time and communication required to privately query a central database of 1,000,000 entries for CODIS systems deployed at different countries (see Online Methods). Here, the client and server are Amazon EC2 instances, with the client located on the West Coast of the U.S. and the server located on the East Coast of the U.S.

| CODIS System | End-to-End Time (seconds) | Bandwidth (MB) |
| --- | --- | --- |
| US System, current (20 STRs) | 38.4 | 172.4 |
| US System, pre-2017 (13 STRs) | 27.7 | 114.9 |
| UK-like System (11 STRs) | 23.4 | 97.3 |
| EU-like System (16 STRs) | 34.1 | 154.5 |
| Chinese-like System (20 STRs) | 40.0 | 189.0 |

By definition, the threshold matching function TM_*k*_ can now be expressed as

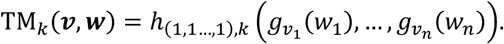

We note that while we can construct a *single* DFA that combines both functionalities, decomposing the computation into two separate steps enables a more efficient privacy-preserving protocol.

We now show how to construct simple DFAs for computing the functions *g*_*v*_ and 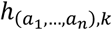. For both functions, we design a “layered” DFA, which can be more efficiently computed in a privacy-preserving manner. First, a DFA consist of a tuple *M* = (*Q*, Σ, *δ, q*_0_, *S*), where *Q* denote the set of states, Σ is the alphabet, *δ*: *Q* × Σ → *Q* is the state-transition function, *q*_0_ ∈ *Q* is the start state, and *S* ⊆ *Q* is the set of accepting states. On input *x* = *x*_1_*x*_2_ ⋯ *x*_*n*_ ∈ Σ^*n*^, the output *M*(*x*) is 1 if *δ*(*q*_*n*−1_, *x*_*n*_) ∈ *S* where *q*_*i*_ = *δ*(*q*_*i*−1_, *x*_*i*_) for all 1 ≤ *i* < *n*, and 0 otherwise. Finally, we say that *M* = (*Q*, Σ, *δ, q*_0_, *S*) is a layered DFA if the following properties hold:

- The set of states *Q* can be partitioned into 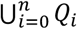 where *Q*_0_, *…, Q*_*n*_ are pairwise disjoint, *Q*_0_ = {*q*_0_}, and *S* ⊆ *Q*_*n*_.
- The state-transition function *δ* can be decomposed into a collection of functions (*δ*_1_, *…, δ*_*n*_) where *δ*_*i*_: *Q*_*i*−1_ × Σ → *Q*_*i*_ and *δ*(*q, σ*) = *δ*_*i*_(*q, σ*) for all 1 ≤ *i* ≤ *n, q* ∈ *Q*_*i*−1_, *σ* ∈ Σ.

In words, a layered DFA is one whose states can be partitioned into a collection of *n* + 1 pairwise disjoint sets (i.e., “layers”) *Q*_0_, *…, Q*_*n*_. On input *x* ∈ Σ^*n*^, the state of the DFA after reading the first *i* bits of *x* is contained in layer *i* (i.e., in the set *Q*_*i*_). We now describe how to represent *g*_*v*_ and 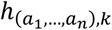 as layered DFAs:

- Let *g*_*v*_(*w*) denote the function that outputs 1 if *v* = *w* and 0 otherwise, where *v, w* ∈ {0,1}^*𝓁*^. It is easy to construct a layered DFA for the equality function. The DFA consists of two branches, each with *𝓁* states: an “accept” branch that corresponds to a matching input and a “reject” branch that corresponds to a non-matching input. Evaluation begins on the accept branch and successively compares the bits of *w* to the bits of *v* (encoded in the DFA transitions). If a mismatch is encountered, the DFA transitions to the reject branch. The output is 1 if the final state is on the accept branch and 0 otherwise. We illustrate this in Figure 1. Thus, for *v* ∈ {0,1}^*𝓁*^, the function *g*_*v*_ can be computed by a layered DFA with *𝓁* + 1 layers, each containing up to 2 states. Note that the vector *v* is entirely encoded in the transition function *δ* of the DFA. Our protocol for privacy-preserving layered DFA evaluation assumes that the topology of *M* is public, but will hide the transition function *δ* (and thus, the value of *v*) from the client.
- Let 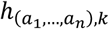 (*b*_1_, *…, b*_*n*_) be the threshold function that outputs 1 if there are at most *k* indices 1 ≤ *i* ≤ *n* where *a*_*i*_ ≠ *b*_*i*_ and 0 otherwise. It is also straightforward to construct a layered DFA for 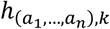. Namely, we consider a DFA with *k* + 2 branches. We label each branch with an integer *j* = 0,1 *…, k, k* + 1, corresponding to the number of mismatches encountered thus far. Evaluation begins on branch 0 and whenever the DFA reads in an input bit *b*_*i*_ ≠ *a*_*i*_, then the state transitions from branch *j* to branch *j* + 1; otherwise, evaluation continues on branch *j*. All state transitions on the final branch *j* = *k* + 1 (corresponding to an input that differs on more than *k* indices), remain on the branch irrespective of the input bit. We illustrate this in Supplementary Figure 2 (for the case where *k* = 1). Thus, the function 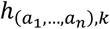 can be implemented by a layered DFA with *n* + 1 layers, each containing up to *k* + 2 states. Note that the sequence of bits (*a*_1_, *…, a*_*n*_) is entirely encoded in the transition function *δ* of the DFA. The topology of the DFA is a function of the dimension *n* and the threshold *k*, both of which will be assumed to be publicly known in our protocol; our final protocol will require that the target bits (*a*_1_, *…, a*_*n*_) be hidden, which holds as long as the protocol for layered DFA evaluation hides *δ*.

It is easy to see that the two layered DFAs described above can be combined into a single layered DFA for computing TM_*k*_. However, the number of layers in the resulting DFA will be the *product* of the number of layers in the two underlying DFAs. When developing our cryptographic protocol for privacy-preserving layered DFA evaluation, both the round complexity and the communication complexity scales with the number of layers. Thus, privately-evaluating two smaller DFAs yields a significantly more efficient protocol compared with evaluating a single larger layered DFA.

### Privacy-preserving evaluation of a layered DFA

We now describe how to use oblivious transfer to construct a privacy-preserving protocol for computing TM_*k*_. We leverage our implementation of the threshold matching function as a *layered* DFA to enable a more efficient protocol. While there already exist protocols for private evaluation of general (i.e., not necessarily layered) DFAs ^32,33^, in this work, we show that the layered structure of our DFAs are amenable to a simpler and direct OT-based evaluation procedure (without needing to additionally rely on heavier cryptographic primitives such as homomorphic encryption).

Let *M* = (*Q*, Σ, *δ, q*_0_, *S*) be a layered DFA: namely, 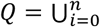 *Q*_*i*_ and *δ* = (*δ*_1_, *…, δ*_*n*_). We will label the states *Q*_*i*_ in layer *i* by indices 1,2, *…*, |*Q*_*i*_|. In our model, the server holds the layered DFA *M* while the client holds the input *x* ∈ Σ^*n*^. At the end of the protocol, the server should learn nothing while the client should learn the value *M*(*x*). In addition, we assume that the *topology* of the DFA *M* is publicly-known (i.e., the client knows the number of layers in *M* as well as the number of states in each layer of *M*). What is hidden is the transition function *δ*. This assumption is true for the setting we consider in this work: here, the CODIS algorithm itself is public (and this determines the topology of the DFA), but the records within the server’s database determine the exact transition function, which is precisely what our protocol hides.

Our protocol relies on the following simple observation: the transition function *δ*_*i*_ can be described by a truth table of size |*Q*_*i*_| ⋅ |Σ|. This yields the following general approach for evaluating *M* privately:

- The client initializes her state to *s*_0_ ← *q*_0_ ∈ *Q*_0_.
- Let *x* ∈ Σ^*n*^ be the client’s input. We maintain the following invariant. At the beginning of round *i* = 1, *…, n* − 1, the client knows the state *s*_*i*−1_ ∈ *Q*_*i*−1_ of the DFA after reading the prefix *x*_1_ ⋯ *x*_*i*−1_. The client wants to learn *s*_*i*_ ≔ *δ*_*i*_(*r*_*i*−1_, *x*_*i*_). To do so, the client and server run a 1-out-of-(|*Q*_*i*_||Σ|) OT where the receiver’s input is (*s*_*i*−1_, *x*_*i*_) and the server associates the message *y*_*i*_ = *δ*_*i*_(*q, σ*) with index (*q, σ*) ∈ *Q*_*i*−1_ × Σ is. At the end of this protocol, the client learns the state *s*_*i*_ = *δ*_*i*_ (*s*_*i*−1_, *x*_*i*_) ∈ *Q*_*i*_ of the DFA after reading *x*_1_ ⋯ *x*_*i*_.
- After *n* − 1 rounds, the client learns the state *s*_*n*−1_ ∈ *Q*_*n*−1_. For the final round, the client and server run a 1-out-of-(|*Q*_*n*−1_||Σ|) OT protocol where the client’s input is (*s*_*n*−1_, *x*_*n*_) and the server associates the value 1 with the index (*q, σ*) ∈ *Q*_*n*−1_ × Σ if *δ*_*n*_(*q, σ*) ∈ *S* (i.e., *δ*_*n*−1_(*q, σ*) is an accepting state) and value 0 for the remaining indices (*q, σ*) where *δ*_*n*_(*q, σ*) ∉ *S*.

Correctness of the above protocol follows by correctness of the underlying OT protocol. In particular, OT is used to iteratively evaluate the layered transitioned functions *δ*_1_, *…, δ*_*n*_. Moreover, privacy for the client’s input is ensured by security of the OT protocol since the server’s view in the entire protocol execution only consists of its view in the OT queries (which hide the client’s input).

However, this protocol does *not* provide privacy for the server. Namely, the client learns the sequence of states corresponding in the DFA evaluation on her input, which could reveal information about the state-transition functions. Consider for instance a layered DFA with *k* branches where the node in branch *j* of layer *i* is always labeled with the index *j*. Then, the sequence of states the client obtains by executing the above protocol completely reveals which branch of the computation her input takes at each step in the DFA evaluation. This in turns leaks information about the transition function *δ*_*i*_ (e.g., the client learns whether *δ*_*i*_(*s*_*i*−1_, *σ*) outputs a state in the *same* branch or in a *different* branch). In the case of our threshold matching DFA, this in turn reveals to the client partial matches between her query and the server’s database entry.

The above attack shows that additional randomization is *necessary* to ensure security for the server. The fix is simple: the server simply *blinds* the index of each state in each round of the protocol. The full protocol is described below:

1. The client initializes her current state to *s*_0_ ← 1. The server initializes *α*_0_ ← 0.
2. On round *i* = 1, *…, n* − 1 of the protocol, the client makes an OT query on the index (*s*_*i*−1_, *x*_*i*_). On round *i* of the protocol, the server samples a random blinding factor *α*_*i*_ ← {1, *…*, |*Q*_*i*_|}. It prepares a table of size |*Q*_*i*_| ⋅ |Σ| where the message *y*_*i*_ associated with index (*q, σ*) ∈ *Q*_*i*−1_ × Σ is *y*_*i*_ = *δ*_*i*_(*q* − *α*_*i*−1_ Mod |*Q*_*i*−1_|, *σ*) + *α*_*i*_ Mod |*Q*_*i*_|. The server uses this collection of |*Q*_*i*_| ⋅ |Σ| as its set of messages in the OT protocol. Let *s*_*i*_ ∈ {1, *…*, |*Q*_*i*_|} be the client’s output in the OT protocol.
3. On the final round of the protocol, the client and server run a 1-out-of-(|*Q*_*n*−1_||Σ|) protocol where the client’s input is (*s*_*n*−1_, *x*_*n*_) and the server associates the message 1 with index (*q, σ*) ∈ *Q*_*n*−1_ × Σ if *δ*_*n*_(*q* − *α*_*n*−1_ mod |*Q*_*n*−1_|, *σ*) ∈ *S* and 0 otherwise.

#### Correctness

Let *z*_0_ = 1, *z*_1_, *…, z*_*n*_ be the sequence of states in the evaluation of *M*(*x*). Namely, *z*_*i*_ = *δ*_*i*_(*z*_*i*−1_, *x*_*i*_) for each 1 ≤ *i* ≤ *n*. First, we show that for each 0 ≤ *i* ≤ *n, s*_*i*_ − *α*_*i*_ = *z*_*i*_ Mod |*Q*_*i*_|. We proceed inductively. The claim trivially holds for *i* = 0. By correctness of the OT protocol,

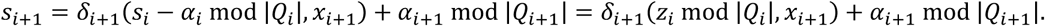

Thus,

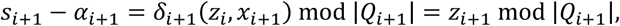

and the claim holds. In the final OT, the output is 1 if

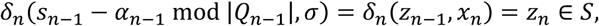

and 0 otherwise. Similarly, the output of *M*(*x*) is 1 if and only if *z*_*n*_ ∈ *S*, and correctness follows.

#### Security

We show that in the OT-hybrid model, the above protocol is secure in the presence of semi-honest adversaries. Recall that in the OT-hybrid model, the client and server interact with an “ideal” OT functionality (i.e., the client sends an index *i* ∈ {1, *…, k*} to the OT functionality, the server sends a collection of messages *m*_1_, *…, m*_*k*_, and the OT functionality replies to the client with *m*_*i*_). If we then instantiate the OT with any concrete protocol that is secure against semi-honest adversaries, we obtain a secure protocol for evaluating layered DFAs. We consider client and server security separately:

- **Client Security (Query Privacy).** In the OT-hybrid model, the server only provides inputs to the ideal OT functionality. It does not receive any message from either the client or the ideal OT functionality. Thus, the view of the server is *independent* of the client’s query so security for the client against a semi-honest server is immediate.
- **Server Security (Function Privacy).** In the OT-hybrid model, the client receives a sequence of values *s*_1_, *…, s*_*n*_ from the ideal OT functionality. From the above correctness analysis, we have that *s*_*n*_ = *M*(*x*) ∈ {0,1}. By construction (and correctness of the OT protocol), each of the intermediate indices *s*_*i*_ for 1 ≤ *i* < *n* satisfies *s*_*i*_ = *δ*_*i*_(*s*_*i*−1_, *x*_*i*_) + *α*_*i*_ Mod |*Q*_*i*_|. Here, the server samples each *α*_*i*_ uniformly from the set {1, *…*, |*Q*_*i*_|}, and in particular, independently of *δ*_*i*_. This means that the value of each *s*_*i*_ is uniform and independent over {1, *…*, |*Q*_*i*_|}. (Formally, we can construct a simulator that on input *M*(*x*) outputs random indices *s*_*i*_ ← {1, *…*, |*Q*_*i*_|} for 1 ≤ *i* < *n* and *s*_*n*_ = *M*(*x*). The output of this simulator is identically distributed as the receiver’s view in the protocol. Thus, the protocol provides perfect security in the OT-hybrid model.)

#### Efficiency

For a layered DFA with *n* + 1 layers, the protocol as described requires *n* rounds of communication, where on round 1 ≤ *i* ≤ *n*, the client and server perform a single 1-out-of-(|*Q*_*i*−1_||Σ|) OT on ⌈log|*Q*_*i*_|⌉-bit messages. Using precomputed OT correlation, the total communication in bits is then

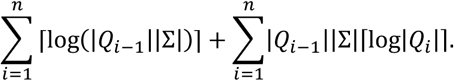

### Secure evaluation of the threshold matching function

We now describe how to use our layered DFAs evaluation protocol to obtain a secure protocol for the threshold matching function TM_*k*_(***v, w***). As discussed above, we can write TM_*k*_(***v, w***) = *h*_(1,1,…,1),*k*_ 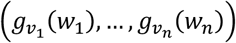 and also represent the functions 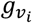 and *h*_(1,1,…,1),k_ as layered DFAs. We can leverage our protocol for secure layered DFA evaluation to evaluate 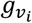 and *h*_(1,1,…,1),*k*_. However, directly applying our privacy-preserving protocol for layered DFAs does not yield a secure protocol for evaluating TM_*k*_ since such a protocol would leak the intermediate values 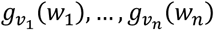 to the client. This means the client would learn the *exact* set of components that match between its vector ***w*** and the server’s vector ***v*** (irrespective of whether the threshold is satisfied or not). We address this by *blinding* the output of the equality-checking circuits.

As currently defined 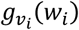 outputs 1 if *v*_*i*_ = *w*_*i*_ and 0 otherwise. We can blind the equality bit by having the server sample a uniform bit *a*_*i*_ ← {0,1} and define 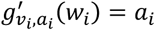 if *v*_i_ = *w*_i_ and 1 – *a*_*i*_ otherwise. We make a few simple observations:

- If we have a layered DFA for computing 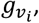, then the same DFA (from Figure 1) can be used to compute 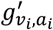 by swapping the accept and reject states in the final layer of the DFA.
- Let 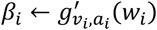. By construction, this means that 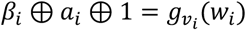. If the client and server apply the privacy-preserving protocol for evaluating a layered DFA to 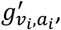, then at the end of the protocol, the client learns *β*_*i*_ (and nothing more) while the server learns nothing. In this case, *β*_*i*_ is uniformly random (it is perfectly hidden by *a*_*i*_), and thus, the client does not learn anything about the value of 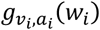. (More precisely, we say that the client and the server have a “secret sharing” of the negated equality bit 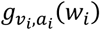 ⊕ 1.)
- The client and server use the privacy-preserving protocol for computing layered DFAs to compute the quantity

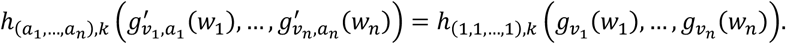

The overall protocol is now given as follows:

1. The server begins by sampling *a*_1_, *…, a*_*n*_ ← {0,1}. For each *i* = 1, *…, n*, the client and server execute the protocol for layered DFA evaluation for the function 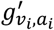. On the *i*^th^ iteration of the protocol, the client provides *w*_*i*_ as its input. Let *b*_1_, *…, b*_*n*_ ∈ {0,1} be the client’s output in the *n* protocol executions.
2. The client and server execute the protocol for layered DFA evaluation for the function 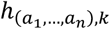 where the client uses *b*_1_, *…, b*_*n*_ as its input. The output *z* is the result of the threshold matching function TM_*k*_(***v, w***).

#### Correctness

Correctness of the above protocol follows via correctness of the underlying layered DFA evaluation protocol. Take any two inputs ***v*** and ***w***. Then, we have that 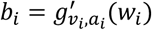 for all 1 ≤ *i* ≤ *n*. The client’s output at the end of the computation is

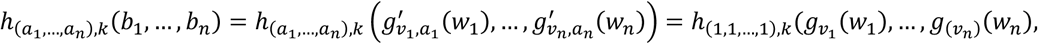

which is precisely the value of TM_*k*_(***v, w***).

#### Security

Security of the protocol also follows by security of the underlying protocol for evaluating layered DFAs. As before, we consider client and server security separately:

- **Client Security (Query Privacy).** The threshold matching protocol consists of *n* + 1 invocations of the protocol for layered DFA evaluation. Security of our protocol for layered DFA evaluation ensures that the server does not learn anything about the client’s input in any of these protocol executions, and security follows. (More formally, security of the layered DFA evaluation protocol implies that the server’s view can be simulated *without* knowledge of the client’s input or output, and correspondingly, the server’s view in the threshold matching protocol can be simulated by simply running the simulator for the underlying layered DFA evaluation protocol).
- **Server Security (Database Privacy).** Security for the server follows similarly from security of the underlying layered DFA evaluation protocol. In the above protocol, the client learns the values of *b*_1_, *…, b*_*n*_ and *z*. From the correctness analysis, we have that *z* = TM_*k*_(***v, w***). Next, by security of the layered DFA evaluation protocol (as applied to the computation of 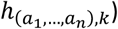), the client learns no additional information other than the value *z*. In particular, the protocol does not leak any information about the values of *a*_1_, *…, a*_*n*_ to the client. By correctness of the layered DFA evaluation protocol, each bit *b*_*i*_ the client obtains satisfies 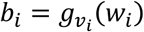 *a*_*i*_⊕ 1. Security of the layered DFA evaluation protocol (as applied to the computation of 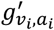) implies that the client learns nothing more about *v*_*i*_, *a*_*i*_ other than the value 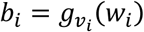 ⊕ *a*_*i*_ ⊕ 1. In particular, this means that the client does not learn anything about the value of *a*_*i*_ from the layered DFA protocol executions. Since the server samples *a*_*i*_ ← {0,1} to be a uniformly random bit, the distribution of *b*_*i*_ is also uniform (even conditioned on the client’s view of the protocol execution, which is independent of *a*_*i*_). As such, the client’s view in the protocol execution can be described by a sequence of uniform random bits *b*_1_, *…, b*_*n*_ together with the output *z* = TM_*k*_(***v, w***). Thus, the client does not learn anything more about *v* other than the value TM_*k*_(***v, w***). (More formally, we can simulate the client’s view of the protocol based on the value of *z* = TM_*k*_(***v, w***) and then sampling the bits *b*_1_, *…, b*_*n*_ ← {0,1}).

#### Efficiency

We now analyze the cost of privately computing TM_*k*_(***v, w***) for two *n*-dimensional vectors ***v, w*** with *n*-bit components where each component *v*_*i*_, *w*_*i*_ ∈ {0,1}^*𝓁*^ is an *𝓁*-bit string.

- Equality checking. Computing the (blinded) equality-check functions 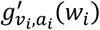 requires evaluating a DFA with *𝓁* + 1 layers, where each layer contains 2 nodes. This requires *𝓁* rounds of communication and 6*𝓁* bits of communication.
- Thresholding. Computing the threshold function 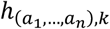 requires evaluating a DFA with *n* + 1 layers, where each layer contains *k* + 2 nodes. This requires *n* rounds of communication and *n*(1 + (2*k* + 5)⌈log(*k* + 2)⌉) bits of communication.

Finally, we note that the equality checks can be conducted in *parallel*. Thus, the round complexity of the complete protocol is *n* + *𝓁* and the total communication (in bits) is

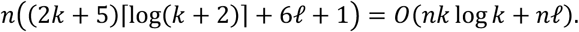

We can achieve a tradeoff between communication complexity and round complexity by having each DFA read multiple bits of the input at a time. This is equivalent to considering a DFA with a larger alphabet (i.e., Σ = {0,1}^*m*^ if the DFA is reading *m* bits of the input for each state transition). Reading multiple bits of the input per state transition decreases the number of layers in the DFA (thus decreasing the round complexity of the private layered DFA evaluation protocol), but increases the size of the transition table between each layer (thus increasing the communication complexity of the protocol). In particular, if we consider DFAs that read *m*-bits of input per transition, then the overall round complexity of the protocol becomes 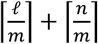 while the communication complexity (in bits) is

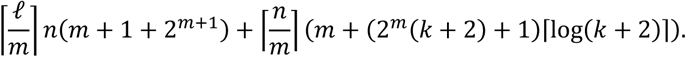

When *𝓁* and *m* are even, then setting *m* = 2 (i.e., reading 2 bits at a time) yields a protocol with smaller round complexity compared to the *m* = 1 setting (by roughly a factor of 2) *and* slightly smaller communication complexity (compared to both the case where *m* = 1 and all *m* > 2). This is the setting we use in our implementation.

### Implementation and evaluation

We implemented the protocol in C++. For our benchmarks, we conducted experiments using two Amazon EC2 instances (M4.2xlarge). Each instance has an 8-core 2.4 GHz Intel Xeon E5-2676 v3 (Haswell) processor and 32 GB of memory. The client instance is located on the West Coast (Northern California region) while the server is located on the East Coast (Northern Virginia region) to simulate a wide-area network (WAN). The network latency between the two instances is roughly 60ms and the bandwidth is roughly 25 MB/s. We use a single-threaded execution environment for all of our experiments.

### Code availability

All code will be made publicly available on Github for non-commercial use upon publication.

## Author Contributions

JAB, KAJ, GB, and DJW designed the study, analyzed results, and wrote the manuscript. JAB wrote software for the analysis with input from KAJ, GB, and DJW.

## Acknowledgements

We thank Benton Case and Dan Boneh for helpful discussions in an early phase of this project and Aviv Regev for support (K.A.J.). This work was also supported by the Joint University Microelectronics Program (JUMP) Undergraduate Research Initiative (J.A.B), the Stanford A.I. Lab (G.B), NSF CNS-1917414 (D.J.W) and a University of Virginia SEAS Research Innovation Award (D.J.W).

---

**Supplementary Figure 1.**
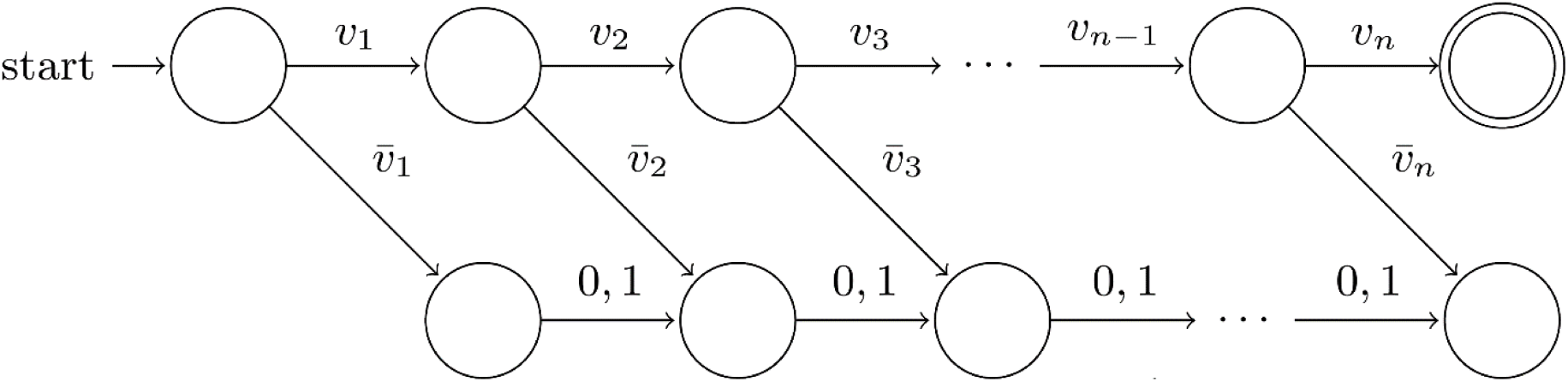
Layered DFA for equality test. This DFA computes the equality-check function *g*_***v***_(***w***) that outputs 1 if ***v*** = ***w*** and 0 otherwise. In particular, for a vector ***v*** = (*v*_1_, *…, v*_*n*_) ∈ {0,1}^*n*^, this DFA only accepts the input ***w*** = (*w*_1_, *…, w*_*n*_) ∈ {0,1}^*n*^ where *v*_*i*_ = *w*_*i*_ for all 1 ≤ *i* ≤ *n*. We use this DFA to decide whether there is a match at a single STR locus. If we denote the single start state as “layer 0”, the two states one can arrive at from layer 0 after reading the first bit as “layer 1”, etc. we see that this DFA has *n* + 1 layers, such that after reading *i* bits, it can only be in one of the two states in layer *i*.

**Supplementary Figure 2.**
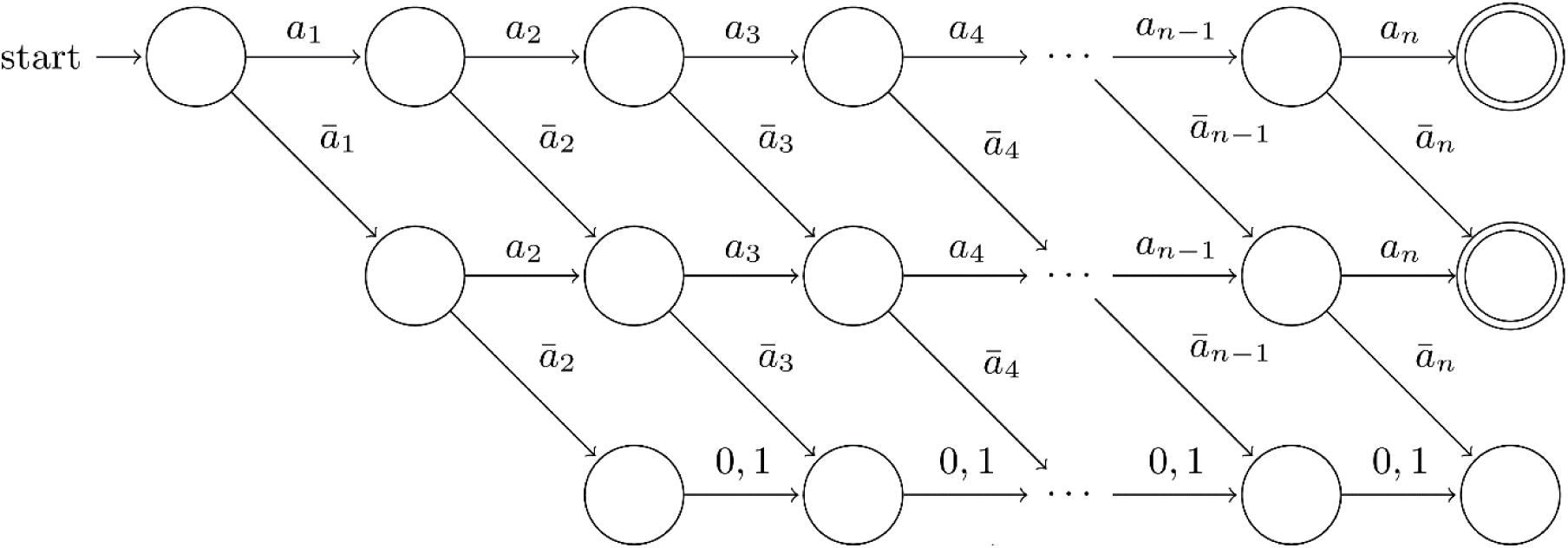
Layered DFA for thresholding. This DFA computes the threshold function 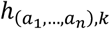 for the *k* = 1 case. Namely, 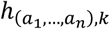(*b*_1_, *…, b*_*n*_) outputs 1 if *a*_*i*_ = *b*_*i*_ for all but at most *k* indices 1 ≤ *i* ≤ *n*. In other words, for any sequence of bits (*a*_1_, *…, a*_*n*_) ∈ {0,1}^*n*^, this DFA accepts if the input *b*_1_, *…, b*_*n*_ satisfies *b*_*i*_ = *a*_*i*_ for all but at most one index *i*. For instance, in this work, we use this DFA to decide whether a DNA profile matches against a database record on at least 19 out of 20 loci (i.e., the setting where *k* = 1 and *n* = 20) as well as the other configurations. Here, the *i*^*th*^ input bit *b*_*i*_ ∈ {0,1} is the (blinded) equality bit denoting whether there is a match in the *i*^*th*^ STR locus (between the agent’s query and the central database’s record). In our protocol, this (blinded) equality bit is computed using the equality-test DFA from Supplementary Figure 1. The bits *a*_1_, *…, a*_*n*_ in the function description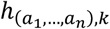 are the blinding values chosen by the server. Recall that the blinding is introduced to hide from the client all information on whether there was a match at STR locus *i* between the database server’s profile and the client’s query. The client only learns whether her query matches the record or not, and nothing more. Much like Supplementary Figure 1, this DFA has *n* + 1 layers, such that after reading *i* bits, the computation can only be in one of the (at most) 3 states of layer *i*.

**Supplementary Table 1.** Approximate CODIS system specifications for select countries. Number of STR loci and number of bits used to encode alleles at each locus for a handful of the 50+ countries using the CODIS system (See Online Methods).

| **CODIS System** | **# of Loci** | **# of Bits** | **Bits per Locus Breakdown** |
| --- | --- | --- | --- |
| **US System, current** | 20 | 212 | 10 bits (17 loci); 14 bits (3 loci) |
| **US System, pre-2017** | 13 | 142 | 10 bits (10 loci); 14 bits (3 loci) |
| **UK-like System** | 11 | 126 | 10 bits (7 loci); 14 bits (4 loci) |
| **EU-like System** | 16 | 178 | 10 bits (12 loci); 14 bits (3 loci); 16 bits (1 locus) |
| **Chinese-like System** | 20 | 224 | 10 bits (14 loci); 14 bits (6 loci) |
| **NIST 40 Loci System** | 40 | 492 | 10 bits (17 loci); 14 bits (23 loci) |

**Supplementary Table 2.** Offline precomputation cost for each setting. Number of oblivious transfer (OT) correlations, and the memory footprint of the OT correlations for the client and server needed to implement a single CODIS search query against a database with 1,000,000 records. Using state-of-the-art OT extension protocols^22^, it is possible to setup 2^24^ > 16 million OT correlations in 7.5 seconds over a wide-area network. For system specifications, see Online Methods.

| CODIS System | Precomputation Size (MB) |  | # of OT Correlations<br>(in millions) |
| --- | --- | --- | --- |
|  | Client | Server |  |
| US System, current (20 STRs) | 122 | 923 | 116 |
| US System, pre-2017 (13 STRs) | 87 | 616 | 78 |
| UK-like System (11 STRs) | 71 | 574 | 73 |
| EU-like System (16 STRs) | 104 | 763 | 96 |
| Chinese-like System (20 STRs) | 122 | 969 | 122 |
| NIST 40 Loci System (40 STRs) | 260 | 2105 | 266 |

